# Covalent protein labeling by SpyTag-SpyCatcher in fixed cells for super-resolution microscopy

**DOI:** 10.1101/125013

**Authors:** Veronica Pessino, Rose Citron, Siyu Feng, Bo Huang

## Abstract

Labeling proteins with high specificity and efficiency is a fundamental prerequisite for microscopic visualization of subcellular protein structures and interactions. While the comparatively small size of epitope tags makes them less perturbative to fusion proteins, they require the use of large antibodies that often limit probe accessibility and effective resolution. Here we use the covalent SpyTag-SpyCatcher system as an epitope-like tag for fluorescent labeling of intracellular proteins in fixed cells for both conventional and super-resolution microscopy. We have also applied this method to endogenous proteins via gene editing, demonstrating its high labeling efficiency and capability for isoform-specific labeling.

---

Epitope tags, short peptides that can be recognized by antibodies, are widely used in fluorescent labeling of fixed-cell proteins for microscopy analysis, especially in the numerous cases where high-quality antibodies directly against the target protein are unavailable. Because of the small size of epitope tags, their use in labeling proteins is less likely to perturb fusion protein function or structural organization compared to using fluorescent proteins or enzymatic tags such as SNAP-tag ^[1]^ and Halo-tag ^[2]^. With recent advancements in genome editing technologies, this small size also facilitates systematic labeling of endogenous genes ^[3]^. However, despite the fact that epitope tags are themselves small, their corresponding antibodies are relatively large. As a result, the staining efficiency is sometimes low due to limited accessibility in tight protein complexes ^[4]^. Additionally, when labeling cellular proteins for super-resolution microscopy, which has been demonstrated to be a powerful approach for dissecting the molecular organization of protein complexes ^[5]^, the size of the antibody is a major concern because it is comparable to the spatial resolution ^[6]^. Although nanobodies (about 1/12 the volume of a full antibody) against larger tags such as GFP have been proven to be an effective approach in reducing probe size ^[7]^, it is inherently difficult to generate high affinity nanobodies for unstructured peptides. Therefore, an epitope tag with a small, tight binder ^[8]^ is highly desirable.

The SpyCatcher-SpyTag system presents a potential solution to many of these challenges. It was engineered by splitting the fibronectin-binding protein (FbaB) of *Streptococcus pyogenes*^[9]^. The SpyTag is a 13 amino acid (a.a.) peptide whose aspartic acid forms a covalent isopeptide bond with a lysine on the 133 a.a. SpyCatcher (Figure 1A). This system has been used in live cells and purified systems to link multiple proteins together ^[10]^. Taking advantage of this tight covalent interaction and the much smaller size of SpyTag compared to antibodies, here we present a method for protein labeling in fixed and permeabilized cells: tagging either over-expressed or endogenous target proteins with SpyTag and then staining with dye-labeled SpyCatcher for fluorescence microscopy, including super-resolution microscopy.

**Figure 1.**
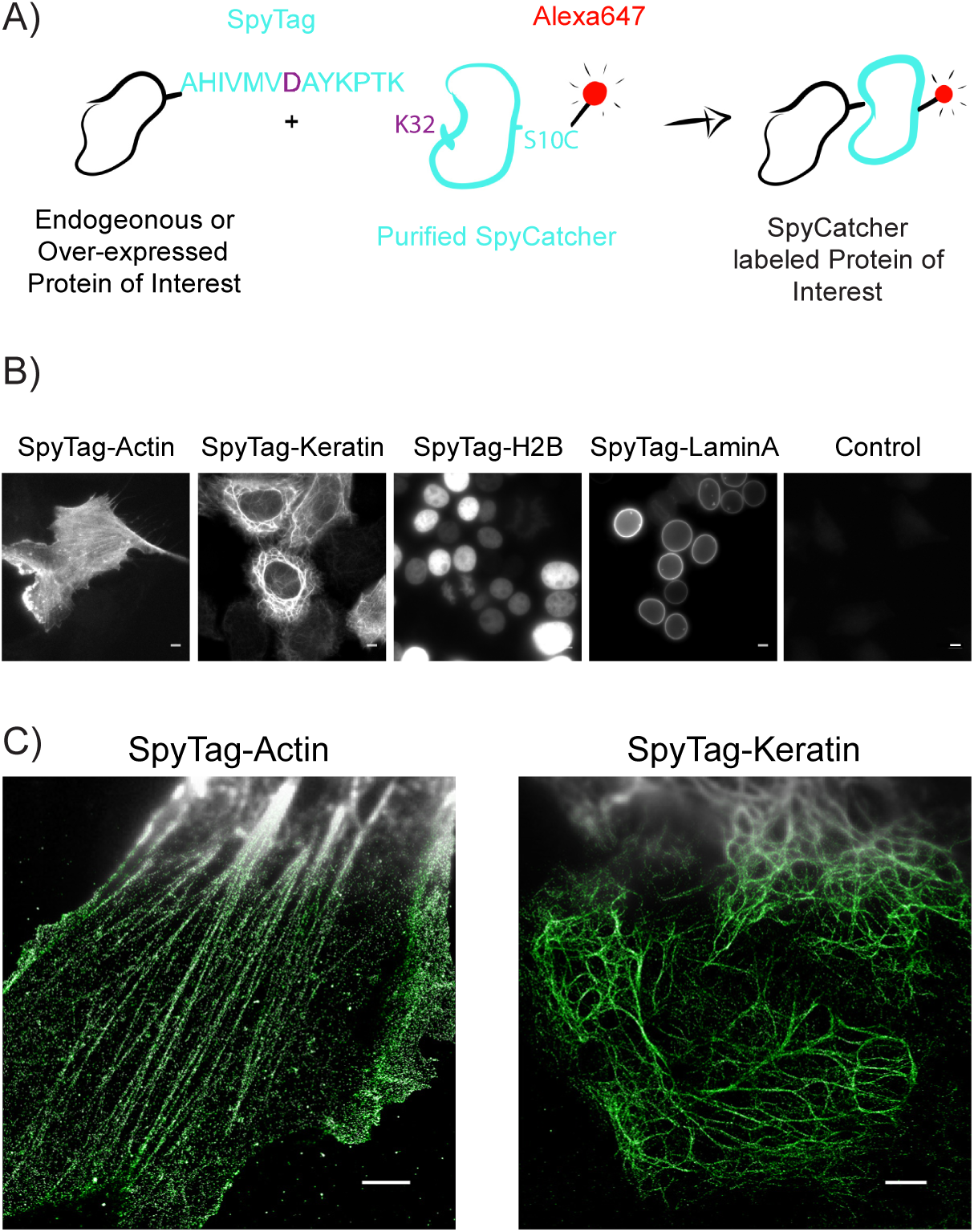
SpyTag-SpyCatcher labelling of proteins in fixed cells. A) A cartoon of the process to use SpyTag as an epitope tag for dye-labelled SpyCatcher in fixed cells. B) Wide-field images of cells stained with SpyCatcher-A647. From left to right: SpyTag-actin overexpressed in HEK293T cells, SpyTag-keratin overexpressed in HeLa cells, SpyTag-H2B overexpressed in HEK293T cells, SpyTag-laminA overexpressed in HEK293T cells, control HeLa cells, no SpyTag expressed. Scale bars: 5 μm. C) Compound wide-field and STORM images of SpyTag-actin overexpressed in RPE-1 cells (left), and SpyTag-keratin overexpressed in HeLa cells (right). Scale bars: 3 μm.

Previously, SpyCatcher and SpyTag have only been used to label extracellular protein loops in live cells ^[11]^. To examine whether SpyCatcher can function intracellularly in fixed and permeabilized cells, in a manner similar to classic antibody staining of epitope tags, we over-expressed SpyTag-actin fusion protein in human retinal pigment epithelial cell line RPE-1 and SpyTag-keratin fusion protein in HeLa cells. Meanwhile, to label the cysteine-free SpyCatcher with a fluorescent dye, we mutated serine-10 to cysteine, purified this recombinant protein and labeled it with Alexa Fluor 647 maleimide (Figure 1A). This labeling scheme ensures a 1:1 dye-to-protein ratio, which benefits super-resolution microscopy using single-molecule switching and localization (commonly known as STORM or PALM) by avoiding the adverse effects of dye-dye interference from over-labeled far-red cyanine dyes ^[6]^. This stoichiometric labelling holds additional potential for quantitative analysis of molecule copy numbers ^[12]^.

After fixing the cells by 2% paraformaldehyde, blocking by bovine serum albumin, and permeabilizing with NP40, over-night staining of SpyCatcher-A647 gave high signal in wide-field fluorescence images of actin and keratin cytoskeletal filaments (Figure 1B). In contrast, negative control cells without transfection displayed negligible background staining. The samples were then imaged with STORM, which clearly resolves actin and keratin fibers that are overlapped in conventional wide-field fluorescence images (Figure 1C). These results show that the SpyCatcher can label intracellular SpyTag specifically and efficiently after fixation.

To demonstrate that the SpyCatcher-SpyTag system has sufficient efficiency to detect proteins expressed at endogenous levels, we took advantage of our previously described method for systematic knock-in of short DNA inserts into endogenous genes ^[3]^. Briefly, we employ CRISPR-Cas9 mediated homology-directed repair to introduce a short synthetic single-stranded DNA oligo into a host cell genome. We tested three target genes: Sec61B, Rab11 and clathrin light chain A (CLTA) (Figure 2A). In all three cases, we were able to obtain high quality wide-field fluorescence images of the endogenous proteins. In particular, the case of Rab11, which does not have highly effective antibodies for immunofluorescence staining, demonstrates the usefulness of our method to label endogenous proteins.

**Figure 2.**
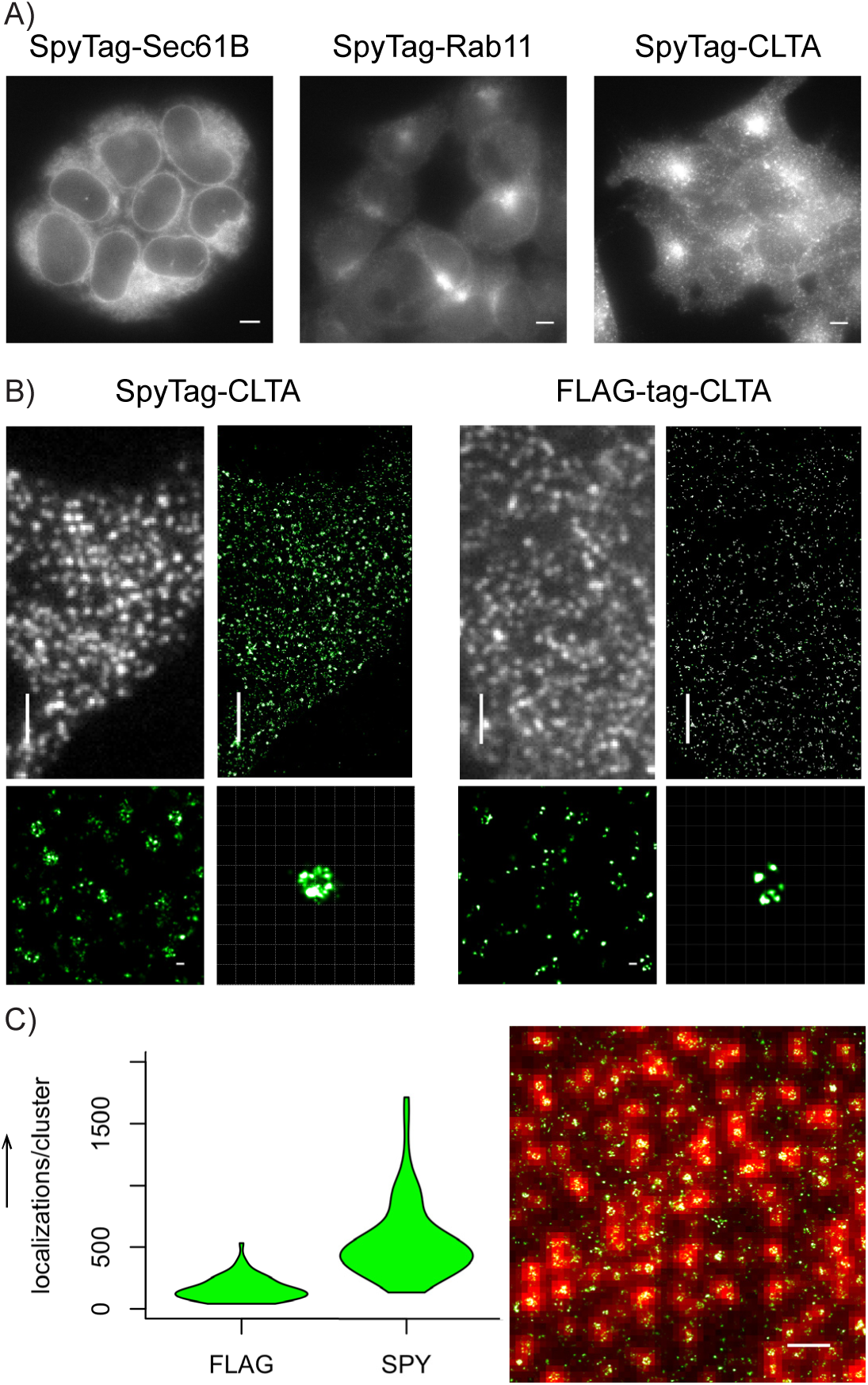
SpyTag-SpyCatcher labelling of endogenous proteins. (A) Wide-field images of knock-in HEK293T cells stained with SpyCatcher-A647: SpyTag-sec61B (left), SpyTag-rab11 (center), SpyTag-CLTA (right). Scale bars: 5 μm. (B) Comparison of wide-field and STORM images of endogenous CLTA labelled by SpyTag knock-in and SpyCatcher staining (left), and by FLAG-tag knock-in cell and anti-FLAG-tag indirect immunofluorescence. Scale bars and grids: 3 μm for wide-field and zoomed-out STORM images, 100 nm for zoomed inserts. (C) Quantification of number of localization points per clathrin structure (*n* = 72 clusters in either condition). Right panel shows an example STORM image overlaid on its corresponding wide-field image used for identifying clathrin structures. Scale bar: 1 μm.

For the case of CLTA knock-in, we compared its labeling using SpyTag and the widely used FLAG-tag. The two tags were knocked in to the same genetic locus and stained by SpyCatcher-A647 and anti-FLAG M2 antibody (then Alexa Fluor 647 labeled secondary antibody), respectively. SpyTag-CLTA cells produced STORM images that clearly revealed the shape of clathrin-coated pits (Figure 2B). Indeed, comparing the A647 signal with the signal from the co-knocked-in GFP marker suggest nearly complete labelling of SpyTag by SpyCatcher, although more careful controls and calibrations are needed to provide an accurate measurement of SpyCatcher labelling efficiency. FLAG-CLTA cells, on the other hand, generated substantially weaker signal under identical acquisition parameters (Figure 2B), with their STORM images containing fewer localization points per clathrin cluster (median: SpyTag = 467, FLAG-tag = 148) (Figure 2C). Consequently, the structural details of clathrin clusters became more difficult to discern. We note that clathrin light chain has two isoforms, CLTA and CLTB, which have mostly interchangeable functionalities ^[13]^. Our labeling of CLTA but not CLTB likely accounted for the sparser labeling of clathrin within a pit by SpyCatcher, compared to previous immunofluorescence images using anti-clathrin light chain antibodies ^[6]^. This difference actually demonstrates the advantage of our approach in distinguishing isoforms, of which many commercial antibodies are not capable.

Altogether, we have demonstrated an efficient and highly specific endogenous protein labeling technique. The use of SpyTag as an epitope for labeling allows for protein visualization with the combined benefits of specificity, covalent bonds, small size, universality and organic dye brightness. We demonstrate its potential for labeling both over-expressed and endogenous proteins, highlighting its potential for targets that lack good antibodies or are subjected to over-expression artifacts. Due to the covalent nature of SpyTag/SpyCatcher linkage, SpyCatcher would be particularly useful in protocols requiring extensive harsh washes such as fluorescence *in situ* hybridization (FISH). Finally, an orthogonal and related covalent-split-protein system was recently developed: SnoopCatcher and SnoopTag ^[14]^. Unfortunately though, fixation seems to inhibit their binding (data not shown), possibly due to the fact that SnoopCatcher-SnoopTag isopeptide bond formation involves a lysine on the Tag instead of the Catcher, which can react with aldehyde fixatives. Future mutagenesis of this residue could reinstate this interaction, which holds potential for an orthogonal system for dual-color labeling.

## Experimental Section

### Cell Culture and Transfection

Human HEK293T and human HeLa cells were grown in Dulbecco’s modified Eagle medium (DMEM) with high glucose and L-Glutamine (Gibco), supplemented with fetal bovine serum (FBS; 10% (vol/vol)) and penicillin/streptomycin (100 μg/ml; UCSF Cell Culture Facility). Human RPE-1 cells were maintained in DMEM/F12 GlutaMAX-I, with sodium bicarbonate (2.438 g/L) and sodium pyruvate (Gibco), supplemented with FBS (10% (vol/vol)) and penicillin/streptomycin (100 μg/ml; UCSF Cell Culture Facility). All cells were grown at 37 °C and 5% CO_2_ in a humidified incubator.

SpyTag-keratin and SpyTag-actin were transfected at 100 ng DNA/well and 1.5 μl of Lipofectamine-2000 (Invitrogen), into an 8-well Lab-TeK II chambered #1.5 coverglass system (Nalge Nunc International).

### Protein Purification and Labelling

SpyCatcher was cloned into the pET28a expression vector and expressed with a N-terminal six-histidine-tag in bacterial BL21 RIL cells. Cells were grown in LB media, induced with IPTG (1 mM) and harvested by centrifugation. Cells were lysed using an Emulsiflex in NaCl (350 mM), HEPES (50 mM) at pH 6.8, Imidazol (20 mM) and 2-mercaptoethanol (5 mM) lysis buffer. Nickel affinity chromagatography was preformed followed by size exclusion on a Superdex 75 (GE Healthcare) column.

Before use, SpyCatcher protein was buffer exchanged into HEPES (50 mM) at pH 6.8 with NaCl (150 mM), using Zeba Spin Desalting Columns (Thermo). Maleimide-Alexa647 C2 (Thermo) dye was then added at a 1:1 molar ratio and quenched with 5:1 excess 2-mercaptoethanol. Excessive dye was removed by running mixture through another Zeba column.

### Molecular Cloning

The DNAs of keratin, β-actin, H2B and LaminA/C were subcloned from mEmerald or mCherry fusion plasmids (cDNA source: the Michael Davidson Fluorescent Protein Collection at the UCSF Nikon Imaging Center). We performed the following restriction enzyme digestion (amino-acid linker length shown in parentheses for each): keratin (18 a.a): mEmerald sequence between BamHI and NotI (mEmerald-Keratin14-N-18); β-actin (18 a.a.): mEmerald sequence between AgeI and BglII (mEmerald-Actin-C-18); H2B (7 a.a.): mEmerald sequence between BmtI and BglII (mEmerald-H2B-C-18); LaminA/C (15 a.a.): mCherry sequence between BmtI and BglII (mCherry-LaminA/C-C-18). The DNA sequences of SpyTag was directly synthesized (Integrated DNA Technologies) and then ligated with the digested vectors using In-Fusion HD Cloning kit (Clontech).

### Knock-in cell line creation and sorting

All synthetic nucleic acid reagents were purchased from Integrated DNA Technologies (IDT). sgRNAs and Cas9/sgRNA ribonucleoprotein (RNP) complexes were prepared as described previously ^[15]^. In order to simplify the isolation of integrated cells, we introduced GFP_11_ (16 a.a.) ^[16]^ in tandem with SpyTag into cells stably expressing GFP_1-10_. Cells were then sorted with fluorescence-activated cell sorting (FACS) using the GFP signal. We note that in practical applications, GFP can be omitted and SpyTag-positive cell can be identified through the classic clonal selection method.

For the knock-in of SpyTag and GFP_11_, 200-nt homology-directed recombination (HDR) templates were ordered in single-stranded DNA (ssDNA) form as ultramer oligos (IDT) (Supplementary Table 1). Cas9 protein (pMJ915 construct, containing two nuclear localization sequences) was expressed in *E.coli* and purified by the University of California, Berkeley Macrolab following protocols described previously ^[17]^. To increase HDR efficiency, HEK293T cells stably expressing GFP_1-10_ were treated with nocodazole (200 ng/mL; Sigma) for 15 hours before electroporation ^[15]^. Cas9/sgRNA RNP complexes were assembled with Cas9 protein (100 pmol) and sgRNA (130 pmol) just prior to electroporation and combined with HDR template in a final volume of 10 μL. Electroporation was performed on an Amaxa 96-well shuttle Nuleofector device (Lonza) via SF-cell line reagents (Lonza).

Nocodazole-treated HEK293T cells stably expressing GFP_1-10_ were resuspended to 10^4^ cells/μL in SF solution immediately before electroporation. For each sample, 20 μL of cells was added to the 10 μL RNP/template mixture. Cells were quickly electroporated using the CM-130 program and transferred to 12-well plate with pre-warmed media. Electroporated cells were cultured for 5-10 days before FACS selection of positive cells. Cell sorting was done in the 488-GFP channel on a FACSAria II (BD Biosciences) in the Laboratory for Cell Analysis at UCSF.

### SpyCatcher and FLAG Immunostaining

Transfected cells were fixed in 2% paraformaldehyde for 30 minutes at room temperature, 48 hours post-transfection, and washed 3 times with phosphate buffered saline (PBS). They were then blocked and permeabilized with 3% Bovine Serum Albumin (BSA) and 2% NP40 for one hour at room temperature. Samples were left in pre-labelled SpyCatcher-A647 (80 nM) in 3% BSA, overnight at room temperature. Endogenous knock-in samples were prepared in the same way. Because the SpyTag-SpyCatcher interaction is stable once formed, samples benefit from longer labeling time at higher temperatures. Finally, samples were washed 5 times over a span of at least 15 minutes.

Of note, blocking with serums was not as efficient as albumin. Fetal Bovine Serum (FBS) and Donkey Neural Serum (DNS) were both tested along with BSA.

FLAG-CLTA cells were plated, fixed and blocked in the same way. Sample was then incubated with primary antibody Anti-FLAG (M2, Sigma) at 1:200 in PBS overnight. The following morning, sample was washed 3 times with PBS, and then incubated with Alexa 647-conjugated secondary antibody for one hour. Finally, cells were washed 5 times with PBS.

### Wide-field Imaging

Wide-field fluorescence images were acquired on an inverted fluorescence microscope (Nikon, Ti-E) with a 100x 1.45 NA oil immersion objective (CFI Plan Apo λ, Nikon). The custom-built epi-illumination optics (Lumen Dynamics, X-Cite XLED1) provided the illumination light at multiple wavelengths. A quadband dichroic mirror (Chroma, ZT405/488/561/640) reflects the illumination light and transmits the fluorescence light. A quad-band emission filter (Chroma, FF410/504/582/669) was used for fast multi-channel fluorescence detection. Another emission filter (Chroma, ET700/50) was also used in the dark red channel to reduce background signal further. The emission filters are mounted onto a motorized filter-wheel (Sutter Instrument, Lambda 10-B), and a motorized *xy*, piezo *z* microscope stage (ASI) controls the translation of the sample. The images were recorded with a sCMOS camera (Hamamatsu, Orca-Flash4.0).

For a rough estimation of SpyCatcher labelling efficiency the estimation, we acquired epi-fluorescence images of the same CLTA-GFP_11_-SpyTag cells in both the GFP and Alexa 647 channels and then normalized the background-subtracted intensities in the two channels by the excitation laser powers, extinction coefficients of the two fluorophores at the respective excitation wavelengths, quantum efficiencies of the two fluorophores, detection efficiencies (integrated fluorescence emission spectra in the filter transmission window) and camera quantum efficiencies at the emission wavelengths. The normalized intensities showed no significant difference between the GFP and Alexa 647 channels.

### STORM image acquisition and analysis

Super-resolution STORM images were collected using a home-built STORM microscope based on a Nikon Eclipse Ti-U inverted microscope. A 405 nm activation laser (OBIS 405, Coherent), and a 647 nm imaging laser (OBIS 647, Coherent) were aligned, expanded, and focused at the back focal plane of the 1.4 NA 100x oil immersion objective (UPlanSApo, Olympus). Images were recorded with an electron multiplying CCD camera (iXon+ DU897E-C20-BV, Andor), and processed via a home-written software. The OBIS lasers were controlled directly by the computer. A quad-band dichroic mirror (ZT405/488/561/640rpc, Chroma) and a band-pass filter (ET700/75m, Chroma for 647nm) separated the fluorescence emission from the excitation light. Maximum laser power used during STORM measured before the objective was 7 μW for 405 nm (∼0.13 W/cm^2^ at the sample), and 24 mW for 647 nm (∼0.46 kW/cm^2^ at the sample). These images were recorded at a frame rate of 60 Hz, with an EMCCD camera gain of 30. During image acquisition, the axial drift of the microscope stage was stabilized by a home-built focus stabilization system utilizing the reflection of an IR laser off the sample. 30,000 frames were collected per sample. STORM imaging buffer made of 100mM TRIS pH 8.0, glucose (10%), NaCl (10 mM), 2-mercaptoethanol (1% v/v), and GLOX scavenging system (1% v/v), was made fresh every 45 minutes. Reconstruction and analysis of the STORM images was performed on the Insight3 software.

To quantify the number of localization points per clathrin-coated structure using SpyTag or FLAG-tag, individual clathrin clusters were identified in conventional fluorescence as fluorescent spots in the Insight3 software using the same algorithm for single-molecule identification. The number of localization points in the corresponding STORM image was then calculated. STORM localization points at nearby positions in consecutive frames were not grouped for this analysis, and such grouping should not affect the relative comparison here. 72 structures were randomly chosen per condition (across 3 cells). These values were then graphed in violin plots, using code written in R.

## Acknowledgements

We thank the C. Craik lab (University of California, San Francisco) for their gift of the SpyCatcher plasmid used for purification; J. DeRisi lab (University of California, San Francisco) for use of their electroporation equipment; and D. Brown of the B.H. lab, for general experimental advice. This work is supported by the National Institutes of Health Director’s New Innovator Award DP2OD008479 (to B.H.) and R21EB022798 (to B.H., S.F. and V.P.). V.P. is supported by a predoctoral fellowship from the American Heart Association. Y.R.C. acknowledges support by the NSF Graduate Research Fellowship. B.H. acknowledges the support from the Chan Zuckerberg Biohub Investigator Program.

---

**[Supplementary Table 1.**
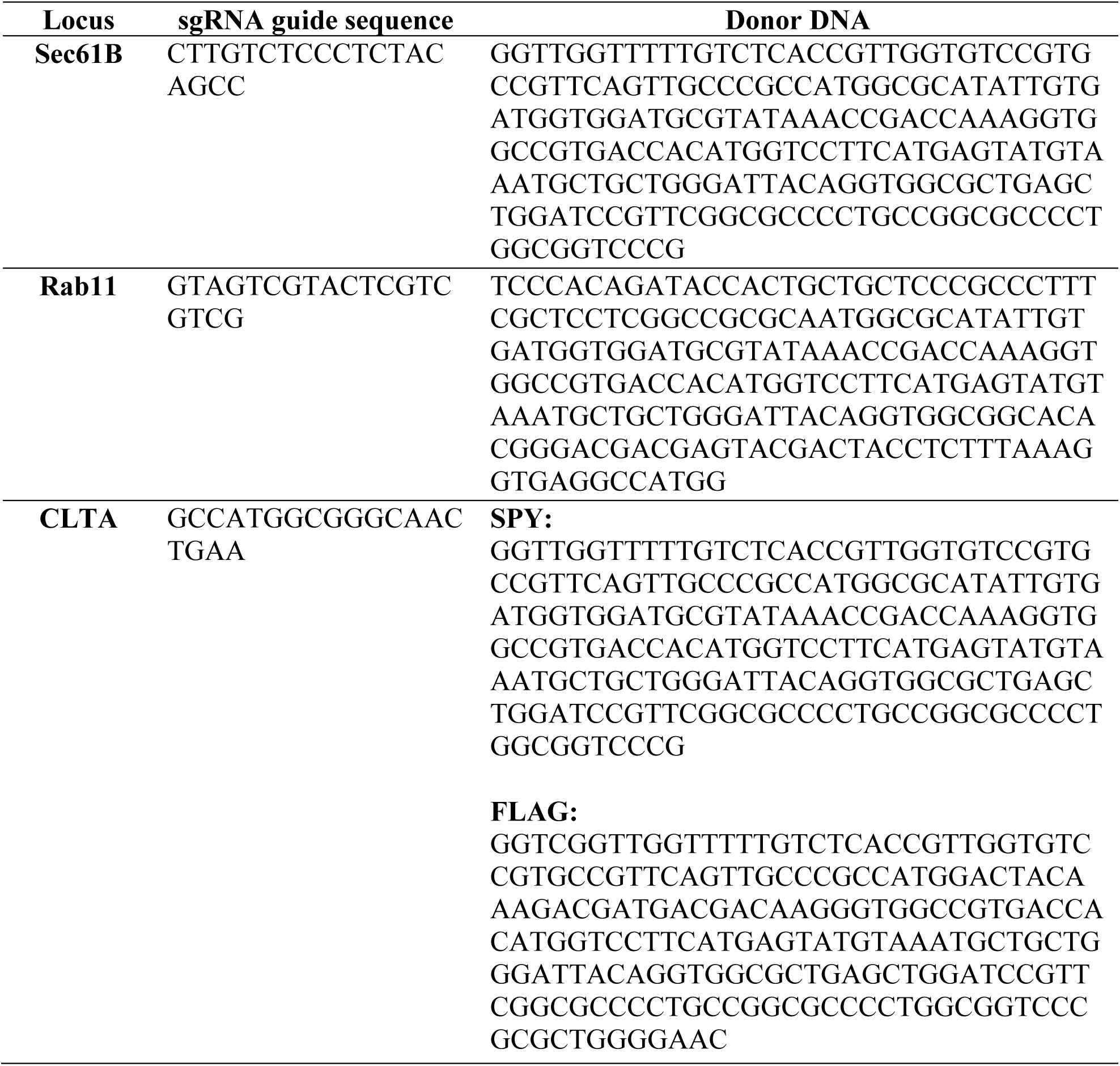
List of DNA sequences used for knock-in cell line creation.

